# Targeting Damaged Collagen for Intra-Articular Delivery of Therapeutics using Collagen Hybridizing Peptides

**DOI:** 10.1101/2022.12.06.519353

**Authors:** E. N. Luke, P. Ratnatilaka Na Bhuket, S. M. Yu, J. A. Weiss

## Abstract

The objective of this study was to investigate the potential of collagen hybridizing peptides (CHPs), which bind to denatured collagen, to extend the retention time of near-infrared fluorophores (NIRF) following intra-articular (IA) injection in rat knee joints. CHPs were synthesized with a NIRF conjugated to N-terminus. Male Sprague-Dawley rats were assigned to one of four experimental groups: healthy, CHP; osteoarthritis (OA), CHP; healthy, scrambled-sequence CHP (sCHP), which has no collagen binding affinity; or OA, sCHP. Animals in the OA groups received an IA injection of monosodium iodoacetate to induce OA. All animals then received the corresponding CHP injection. Animals were imaged repeatedly over two weeks using an *in vivo* fluorescence imaging system. Joint components were isolated and imaged to determine CHP binding distribution. Safranin-O and Fast Green histological staining was performed to confirm the development of OA. CHPs were found to be retained within the joint following IA injection in both healthy and OA animals for the full study period. In contrast, sCHPs were cleared within 24-48 hours. CHP signal was significantly greater (p<0.05) in OA joints when compared to healthy joints. At the two-week end point, multiple joint components retained CHPs, including cartilage, meniscus, and synovium. CHPs extended the retention time of NIRFs following IA injection in healthy and OA knee joints by binding to multiple collagenous tissues in the joint. These results support the pursuit of further research on CHP based therapeutics for IA treatment of OA.

## INTRODUCTION

Treatment options for osteoarthritis (OA) include non-surgical intervention, surgery, and administration of therapeutics, but they each have limitations. Non-surgical interventions, which include physical therapy and lifestyle modifications, are a viable option for early-stage OA, but often do not alleviate pain for patients with more severe OA, which is typically seen in the clinic as pain shows up later in the disease course. Surgical interventions such as arthroscopic revision and total joint replacement are not ideal for all patients. Younger patients are not suitable for joint replacement since artificial joints typically last about 20 years, while older patients have increased risk for complications during surgery and recovery^[1]^. For many decades, locally administered therapeutics have been used to treat OA^[2]^. Local administration of drugs via intra-articular (IA) injection provides site-specific treatment, while circumventing many complications from systemic administration of drugs, including off-target toxicity due to large dosing requirements^[3]^. However, in the highly vascularized joint space, endogenous and exogenous components of synovial fluid are quickly removed through blood and lymphatics^[4,5]^, which reduces the efficacy and duration of activity of the OA drugs intended to alleviate pain or alter disease progression.

Currently, there are no FDA approved disease-modifying OA drugs that provide significant clinical improvements^[6,7]^. In the case of IA injection, rapid clearance of synovial fluid components is a limiting factor. There have been attempts to extend the retention time of therapeutics following IA injection, which include loading drugs into particles of varying size (microparticles and nanoparticles), as well as use of drug delivery system that binds to cartilage surfaces^[8]^. Despite these efforts, retention of even medium size molecules of hundreds of nanometer size remains a major challenge for IA therapeutics because of fast clearance from the joint, especially in the context of increased vascular and lymphatic permeability of inflamed joint tissues. In addition, we have limited understanding of location of IA injected molecules in the joint space and their quantitative clearance profile, particularly in OA joints^[9,10]^. Therefore, there is a critical need to develop a new method to extend the retention time of locally administered joint therapeutics. A promising strategy is the targeting of degraded collagen within the articular cartilage matrix and joint tissue, since collagen makes up ∼60% of the dry weight of cartilage and OA increases collagen degradation in the joint^[11]^.

Collagen hybridizing peptides (CHPs) have the potential to extend retention time of therapeutics in the joint space by binding to damaged collagen. CHPs bind to the unfolded triple helix chains of damaged collagen by formation of triple helix, providing a strong and specific affinity to denatured collagens associated with a wide range of diseases and injury^[11,12]^. We demonstrated this in various collagen types (I, II, III, IV, etc.) using both *in vitro* and *in vivo* models^[11,13–15]^. In particular, we demonstrated that CHPs bind to denatured collagen in mechanically damaged tendons and OA articular cartilage in ex vivo models of positional and energy-storing fascicles and fibrils from rats^[16–20]^, and human and guinea pig articular cartilage^[11,19]^, respectively. CHPs, which have high serum stability^[21]^, have also been shown to bind to many collagenous tissues following systemic injection, with the highest accumulation in the spine and joints which have high collagen remodeling activity^[12,13,21,22]^. Recently, Kiick and coworkers developed self-assembled nanoparticles (∼130 nm) which display collagen like peptides on the surface. They demonstrated more than 7-day retainment of the IA injected nanoparticles in the healthy mouse knee joint. However, due to the lack of a control experiment, it is unclear if this retention was a direct result of the peptide’s affinity to collagen^[23]^.

In this research, we investigated whether CHPs could improve the retention time of small molecules in joint space after IA injection. We examined if CHP conjugation could extend the retention time of near-infrared fluorophores (NIRF) markers *in vivo* after IA injection in healthy and osteoarthritic rat knee joints. Our results demonstrate that CHPs prolong the localization of injected marker molecules in the joint space which represents a significant step towards extending the lifetime of joint therapeutics.

## METHODS

### Materials

A 10 mg/mL solution of monosodium iodoacetate (MIA; Sigma Aldrich, Burlington, MA; CAS: 305-53-3) was prepared in sterile saline. The preparation was made <2 hr before injection to ensure fresh solution was injected^[24]^. NIRF-conjugated peptides were prepared as previously reported with some modifications^[21]^. Briefly, a CHP with a GGG(GPO)_9_ sequence and a scrambled CHP [sCHP; PGO-GPG-POP-OGO-GOP-PGO-OPG-GOO-PPG] were prepared via automated Fmoc-mediated solid phase peptide synthesis using a Focus XC solid phase peptide synthesizer (Aapptec, Louisville, KY, USA)^[21]^. Fmoc-6-Aminohexanoic acid (Fmoc-Ahx) was manually coupled to the N-termini of the CHP and the sCHP by HBTU/HOAt coupling chemistry, followed by the removal of its Fmoc group by 20% piperidine in *N,N*-dimethylformamide. Ahx served as a linker between the peptides and the NIRF dye. The Ahx-attached CHP and sCHP were cleaved from resins and purified by an Agilent Prep-Star high-performance liquid chromatograph (HPLC; Santa Clara, CA, USA). Then, 0.5 mg of IRDye 680RD NHS Ester (Licor, Lincoln, NE, USA) was mixed with 2 mg of the Ahx-attached CHP or sCHP in a 7.5× phosphate-buffered saline (PBS) solution. The conjugation reaction was done overnight at 4 °C under dark condition. The resulting NIRF-CHP and NIRF-sCHP were purified by the HPLC. The purified peptides were then verified by a Bruker maXis-II ETD ESI-qTOF mass spectrometer (Billerica, MA, USA). The purified NIRF-CHP and NIRF-sCHP were lyophilized and subsequently resolubilized in DI water filtered with a 0.22 μm membrane to obtain stock solutions. Then, the NIRF-CHP and NIRF-sCHP stock solutions were separately diluted in sterile saline at a final concentration of 40 μM (1 nmol/25 μL).

### Animals

Male Sprague Dawley rats (N=16) weighing 126-150 g were kept in accordance with the approved IACUC protocol through the University of Utah. In brief, animals were allowed free cage activity, access to food and water ad libitum, and were acclimated for one week prior to experimentation. Animals were randomly assigned to one of four treatment groups using a randomized block design (n=4 per group): healthy, CHP; OA, CHP; healthy, sCHP; or OA, sCHP.

### Induction of OA

Animals were anesthetized using isoflurane (3-5%) and right knee joints were shaved using a Remington beard trimmer. Joints were sterilized with isopropanol and a single 25 μL injection of 10 mg/mL MIA was made into the right knee joint lateral to the patellar tendon^[24–26]^. Joints were flexed and extended ten times to ensure distribution of injected solution throughout the IA space. Animals that received an MIA injection were given 2 mg/kg Meloxicam daily for pain relief starting on the day of OA induction. Animals showed no signs of pain or discomfort during movement, and rearing was comparable to their healthy counterparts.

### CHP Injections and Fluorescence Imaging

OA was allowed to develop for one week prior to injection of the respective CHP solution. Animals were anesthetized using isoflurane (3-5%), right knee joints were shaved again, and skin was sterilized using isopropanol. Both CHP and sCHP solutions were heated at 80 °C for 10 min to activate the peptides followed by quenching on ice for 30 seconds^[11,21]^.

Injections were made within two minutes after CHP quenching. A single IA injection of CHP solution (25 μL) was performed using a 27-gauge, 0.5-inch needle lateral to the patellar tendon. Joints were flexed and extended ten times to ensure distribution of injected solution within the joint space. Animals were kept under anesthesia (3-4%) following CHP injection and an initial image acquisition was performed using an IVIS Spectrum imaging system (PerkinElmer, Waltham, MA) <15 minutes following injection. Fluorescent images were captured with the following parameters: exposure time of 1 sec, medium binning, F/Stop 8, excitation: 675 nm, and emission 720 nm. Field of view “C” (13 cm) and a subject height of 3.5 cm were applied.

Animals were imaged subsequently using the same imaging parameters at 6, 12, 24, 48, 72, 120, 168, 240, and 336 hr following CHP injection. Animals were anesthetized using isoflurane (3-5%) during each imaging s0065ssion.

Regions of interest (ROIs) in the image data were selected to measure the total radiant efficiency 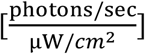 of the NIRF-CHP within the joint and background signal. ROI, which encapsulated the entire joint space, was kept consistent in size between imaging sessions. Background ROIs were also kept consistent in size and centered around the ankle in the shaved region. We accommodated for possible changes in the image acquisition system or parameters between imaging sessions by imaging a well plate containing dilutions of the NIRF-CHP solution. The differences in calibration curves between time points were used to correct total radiant efficiency data, using the calibration curve at the two hr time point as the baseline.

### Tissue Harvesting and Histology

Animals were euthanized by CO_2_ inhalation, followed by exsanguination. Right legs were harvested; briefly, skin and surrounding musculature was removed with joint capsule intact. Legs were fixed in 10% neutral buffered formalin (NBF) for 120 hr at 4 °C. NBF was changed every 24 hr and legs remained on a shaker plate for the period of fixation. Further processing for histological evaluation was performed by Hooke Labs (Lawrence, MA); briefly, joints were decalcified in Immunocal decalcifier (StatLab, Columbia, MD; SKU #: 1414-32), paraffin embedded on a Tissue Tek VIP-5 processor (Sakura Finetek, Torrance, CA), sectioned to 3 μm thickness, and stained using Safranin-O and Fast Green. Histology slides were visualized using an Olympus IX71inverted light microscope with 10× Plan, 20× LCPlanFl, and 40× LUCPlanFl Olympus objectives. The joint space was located and images were taken of the femoral and tibial cartilages with underlying bone. Location in the joint was kept consistent by imaging similar anatomic areas (dorsal to ventral) based on slide number. Images were captured using a Canon 80D camera. Scale bars were added in ImageJ (NIH, Bethesda, MD).

### Isolating Joint Components

Three additional Male Sprague Dawley rats (126-150 g) were used to determine the spatial distribution of CHP binding within the knee joint. Each animal received a single injection of CHPs. After 336 hr, animals were imaged using the IVIS as described above to confirm the presence of CHPs in the joint. Then, animals were euthanized and knee joints were harvested as above. Joints were then further dissected to expose the tibial and femoral cartilage, medial and lateral menisci, and synovial membrane with patella attached. Using a No. 10 scalpel, the joint capsule was opened proximal to the knee joint, followed by cuts on the medial and lateral aspects. The joint capsule, with patella still attached, was folded down towards the tibia and the ligaments stabilizing the joint were cut, taking care not to damage the menisci and cartilage layers. The menisci were carefully isolated and placed in a 1× PBS buffer. Then, excess bone on the femur and tibia were trimmed, allowing for optimal positioning during imaging. Isolated joint components were imaged using the IVIS, with imaging parameters described above, and a Canon 80D digital camera equipped with a 60 mm lens.

### Data Analysis and Statistics

Primary outcomes of this study included differences in retention time between CHP and sCHP, as well as between healthy and OA animals. Secondary outcomes included confirming the development of OA and identifying the distribution of CHP binding in the joint following IA injections. All animals (N=16) were included in the following analyses. Confounding variables, such as order of treatment and animal location, were not controlled.

All data were analyzed in Stata version 16 (College, TX, USA). Sample size was determined using pilot data in the Stata sample size calculator, resulting in a final sample size of n=4 per group. Briefly, total radiant efficiency at each time point compared between the healthy CHP and OA CHP groups and the area under the curve was compared between all groups. The maximum resulting sample size from each of these calculations was used for this study. Data is presented as mean ± 95% confidence interval (CI). The normality of the data was tested using a Shapiro-Wilk test. There was no instance of lack of normality within any groups. Total radiant efficiency vs. time profiles of each rat were analyzed using a noncompartmental model to calculate several pharmacokinetic (PK) parameters including area under the profile curve (*AUC*), total area under the first moment curve (*AUMC*), mean residence time (*MRT*), half-life (*t*_1/2_), and elimination rate constant (*k*_e_). The primary measurement used to test whether CHPs were retained to a greater extent in OA animals compared to healthy animals was AUC, which combined repeated measurements across time into a single statistical summary. We then compared PK parameters between the independent groups using a two sample Student’s t-test. Since these parameters was tested using one significance test, there was no need to adjust for multiple comparisons.

## RESULTS

### Fluorescence imaging of IA injected NIRF-CHPs in OA and healthy rat model

Fluorescence imaging of animals over the two-week period provided information on the retention of both CHP and sCHP following IA injection. Analysis began at the 12-hr mark, when the total radiant efficiency signal stabilized. Fluorescence intensity from *NIRF-CHPs* was clearly observed in the joint space for the entire 336-hr (2 week) imaging period compared to sCHPs which showed negligible intensity by 12-48 hr (Fig. 1). Both the healthy and OA CHP groups had measurable amounts of NIRF-CHP present at the end point, while sCHP signal was completely cleared in healthy and OA joints by 24 and 168 hr, respectively. In OA animals, CHPs were ultimately retained for 2 times longer than sCHPs, whereas in healthy animals, CHPs were retained for 14 times longer than sCHPs. CHPs were retained to a greater extent in the OA animals compared to healthy animals at all imaging time points (p<0.05) (Fig. 2). Mean ± 95% CI calculations confirmed that between group differences were significant for comparisons made between OA and healthy CHP groups (Fig. 3a), OA CHP and OA sCHP groups (Fig. 3b), and healthy CHP and healthy sCHP groups (Fig. 3c).

**Figure 1.**
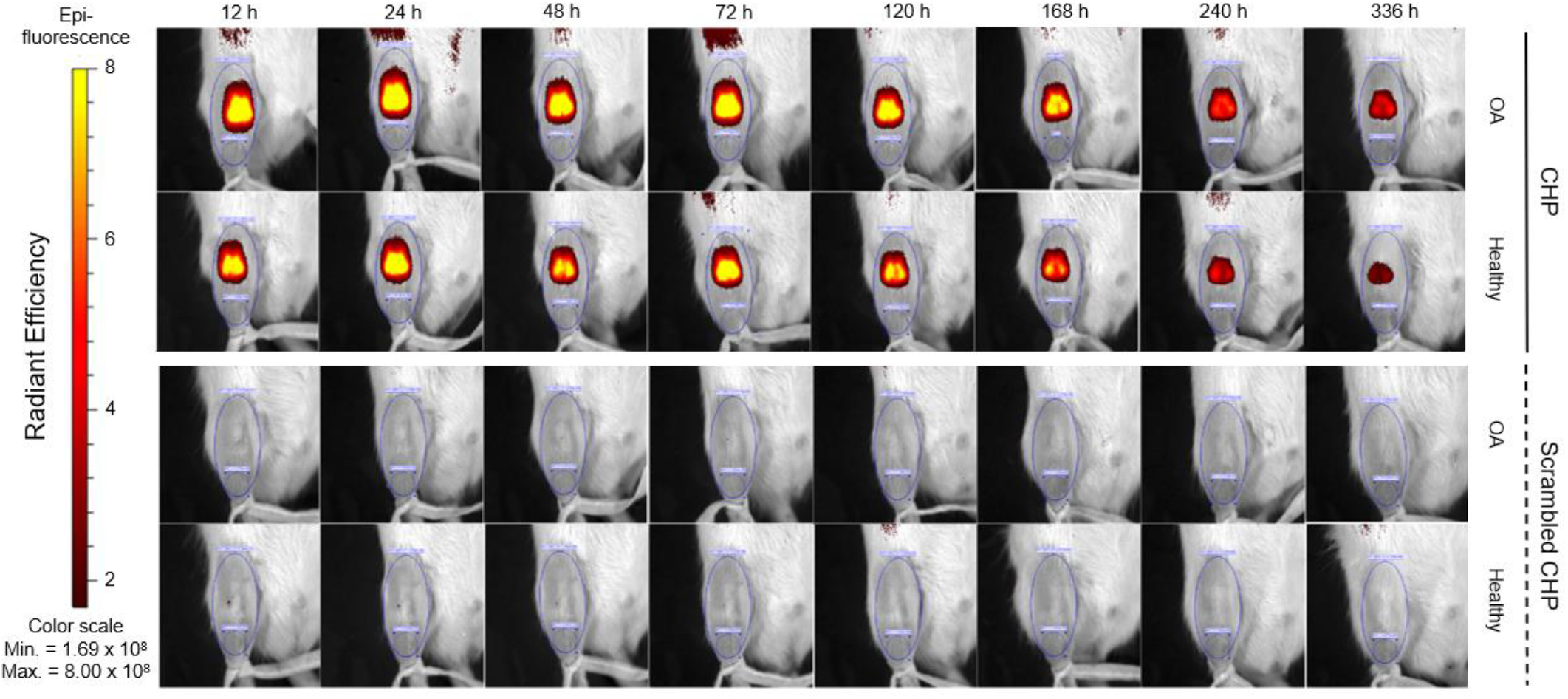
(page width). Time course of retention of NIRF imaging markers in rat stifle joints. Each row shows a single animal in one of the four groups, and each column shows the specific imaging time. In both OA and healthy animals, NIRF-CHPs were retained for the entire imaging period of 336 hrs. In contrast, NIRF-sCHPs in the OA and healthy animals were not visible after 24-48 hours. Representative animal from each group shown (n=4 per group).

**Figure 2.**
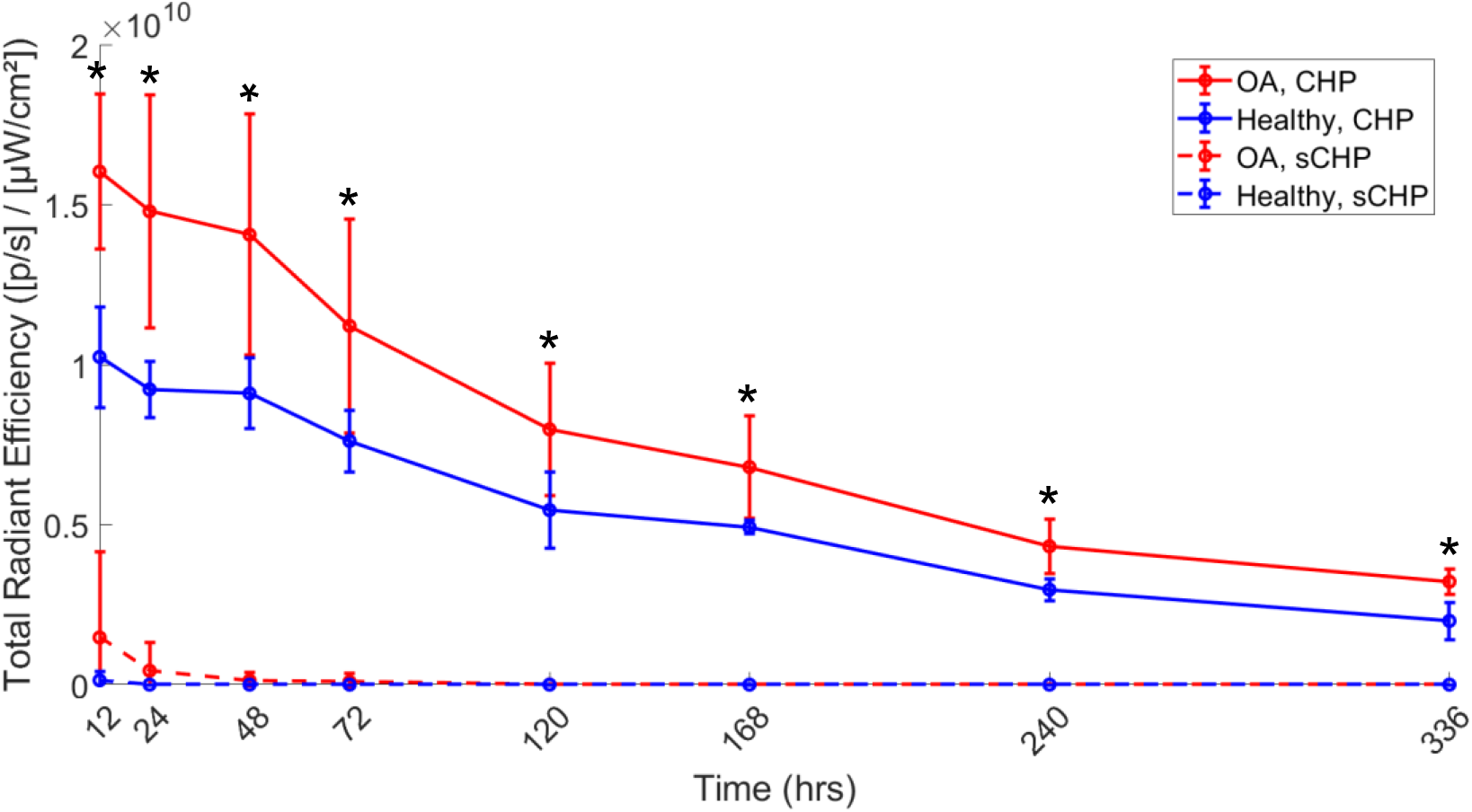
(column width). Total radiant efficiency over the time course of IVIS imaging. CHPs were retained to a greater extent in OA joints (solid red line) compared to healthy joints (solid blue line). sCHPs were cleared from the joint quickly and the total radiant efficiencies were not statistically significant between OA and healthy groups. Data presented as mean ± 95% confidence interval. Asterisk indicates statistical significance between OA CHP and healthy CHP at each time point (p<0.05).

**Figure 3.**
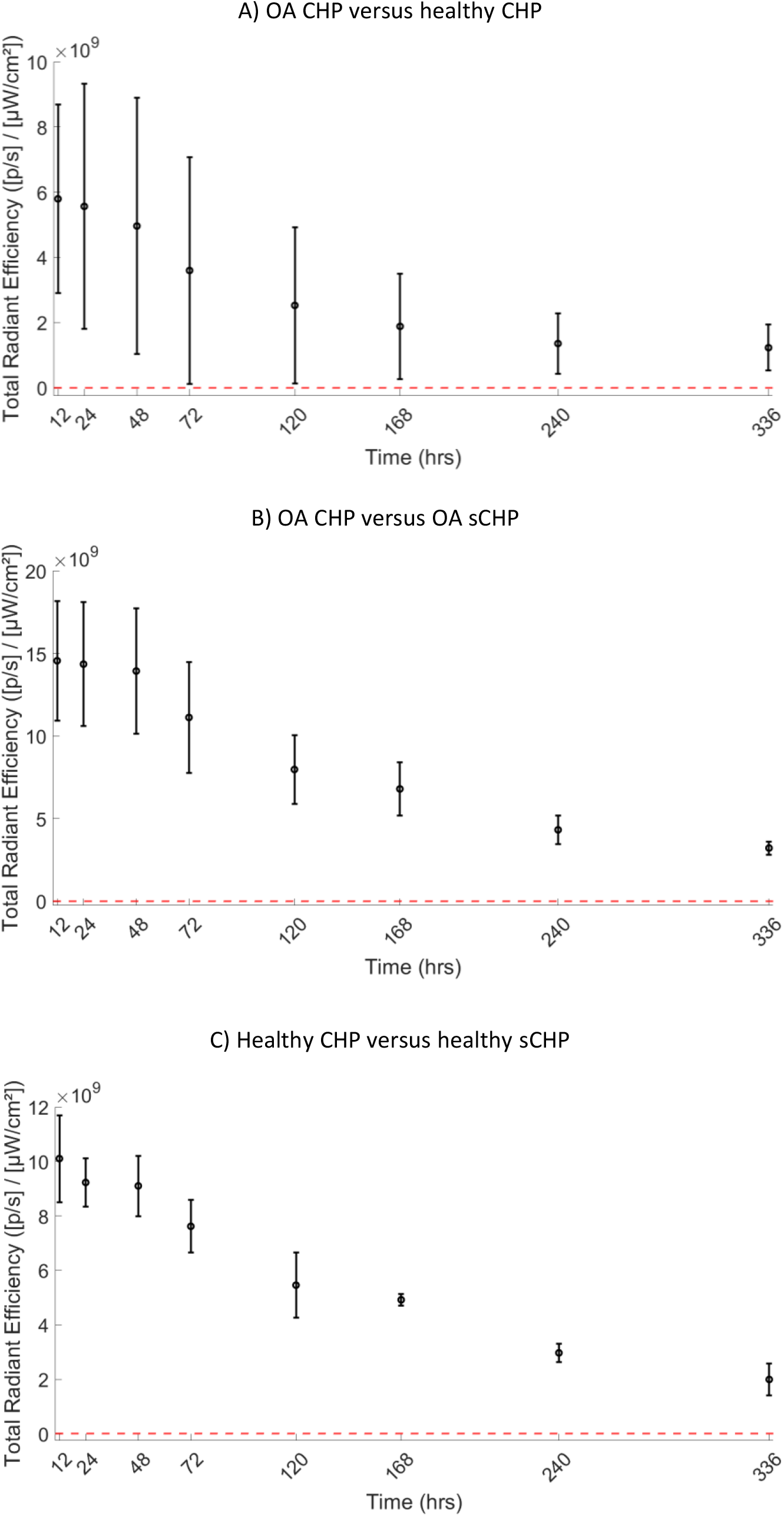
(column width). Differences in total radiant efficiency between group means as a function of time. A) OA CHP versus healthy CHP. B) OA CHP versus OA sCHP. C) healthy CHP and healthy sCHP. Values are represented as mean difference ± 95% CI. There were significant differences in group means for all three comparisons.

We found significant differences in *AUC* for OA CHP compared to OA sCHP (p < 0.0001) and healthy CHP compared to healthy sCHP (p < 0.0001) (Table 1). Interestingly, the OA CHP group showed a significant greater *AUC* compared to the healthy CHP group (p = 0.0015). The *AUMC* of the OA CHP cohort was also significant higher than that of the healthy CHP group (p = 0.0498). These results suggest that the CHP could increase the extent of NIRF retention within the joint spaces, especially in those with pathological lesions. However, there were no significant differences between OA CHP and healthy CHP for *MRT* (p = 0.3414), *t*_1/2_ (p = 0.8697), and *k*_e_ (p = 1.0).

**Table I.**
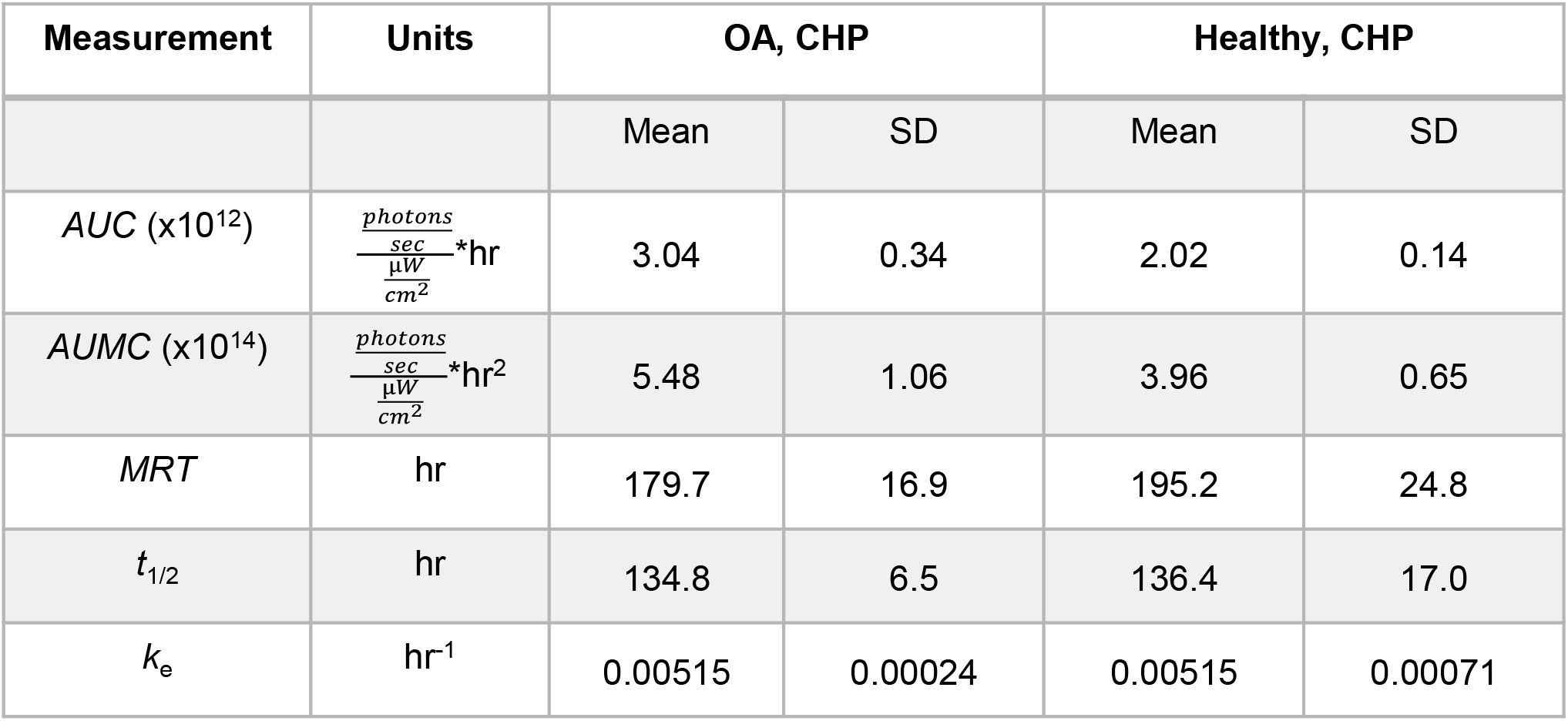
Pharmacokinetic parameters reported for CHP groups (n=4 per group). Area under the curve (*AUC*), total area under the first moment curve (*AUMC*), mean residence time (*MRT*), half-life (*t*_1/2_), and elimination rate constant (*k*_e_) are reported. Values for sCHP groups not calculated due to lack of sufficient data points since sCHP was eliminated from the joint quickly.

### CHP binding distribution in isolated joint components

Following the confirmation of CHP retention 336 hr after IA injection, joint components were isolated to determine the location CHP binding within the IA space (Fig. 4a). Fluorescence imaging revealed CHP staining on all isolated joint components which include cartilages, menisci, and synovial membrane (Fig. 4b). The highest signal intensity was present on the center of the cartilages, which contains the ends of the severed anterior and posterior cruciate ligaments.

**Figure 4.**
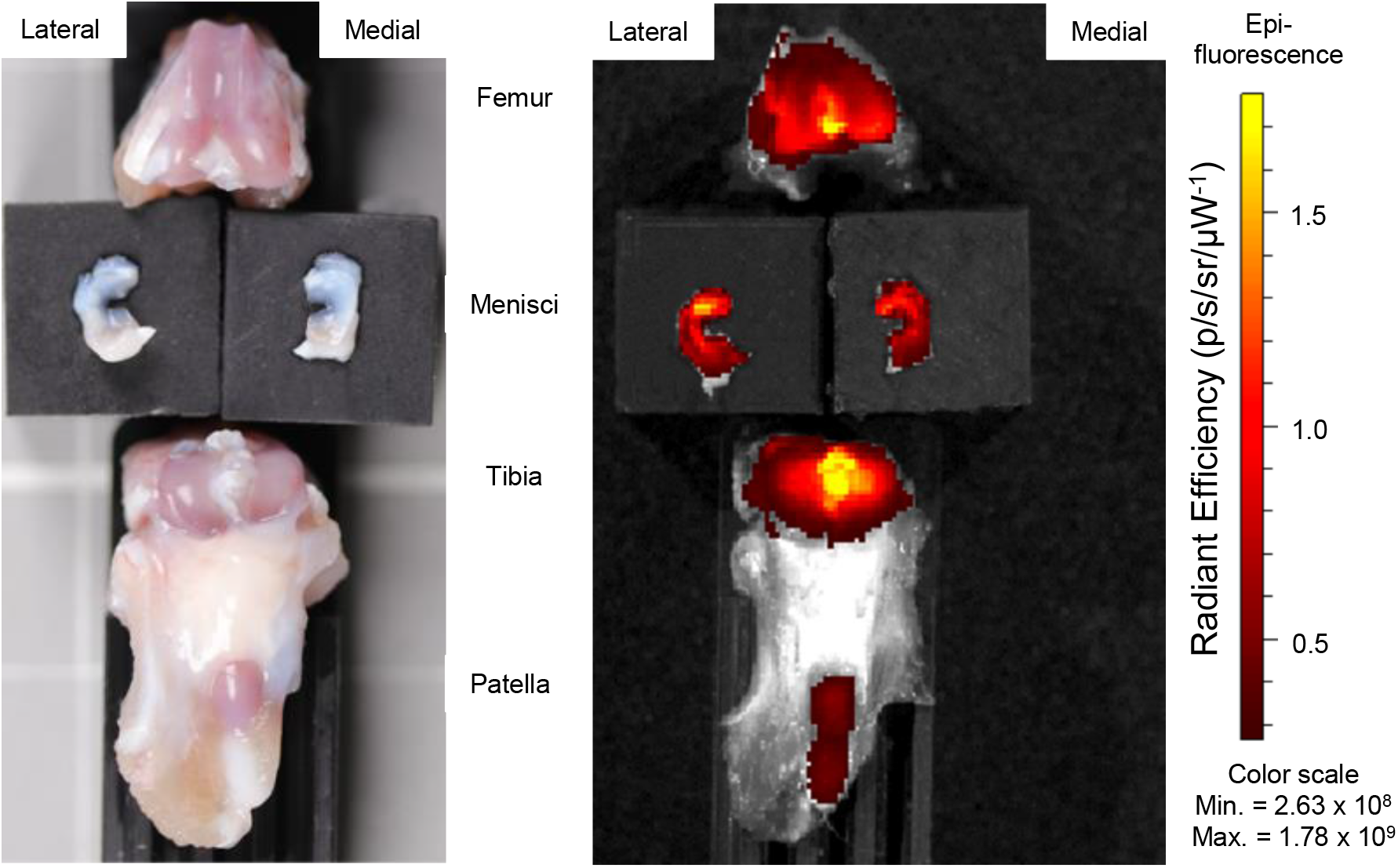
(column width). Healthy dissected stifle joint at 336 hr post-injection of CHPs. A) Macroscopic image of dissected joint components show no apparent disruption to cartilage health following CHP injection. B) IVIS imaging showing binding of CHPs to tibial and femoral cartilage surfaces, medial and lateral menisci, patella, and patellar tendon. The highest signal intensity is localized to the cartilage surfaces.

### Histological evaluation of joint condition in OA and healthy animals

The development of OA in the knee joint was confirmed using histological evaluation. There were no signs of histological change to the cartilage matrix of healthy joints that received CHP injection (Fig. 5a). With the development of OA, there was a noted decrease in proteoglycan staining, loss of chondrocytes, and increase in calcified cartilage at the osteochondral interface (Fig. 5b). Macroscopically, OA was confirmed by the development of lesions on the cartilage surface (data not shown).

**Figure 5.**
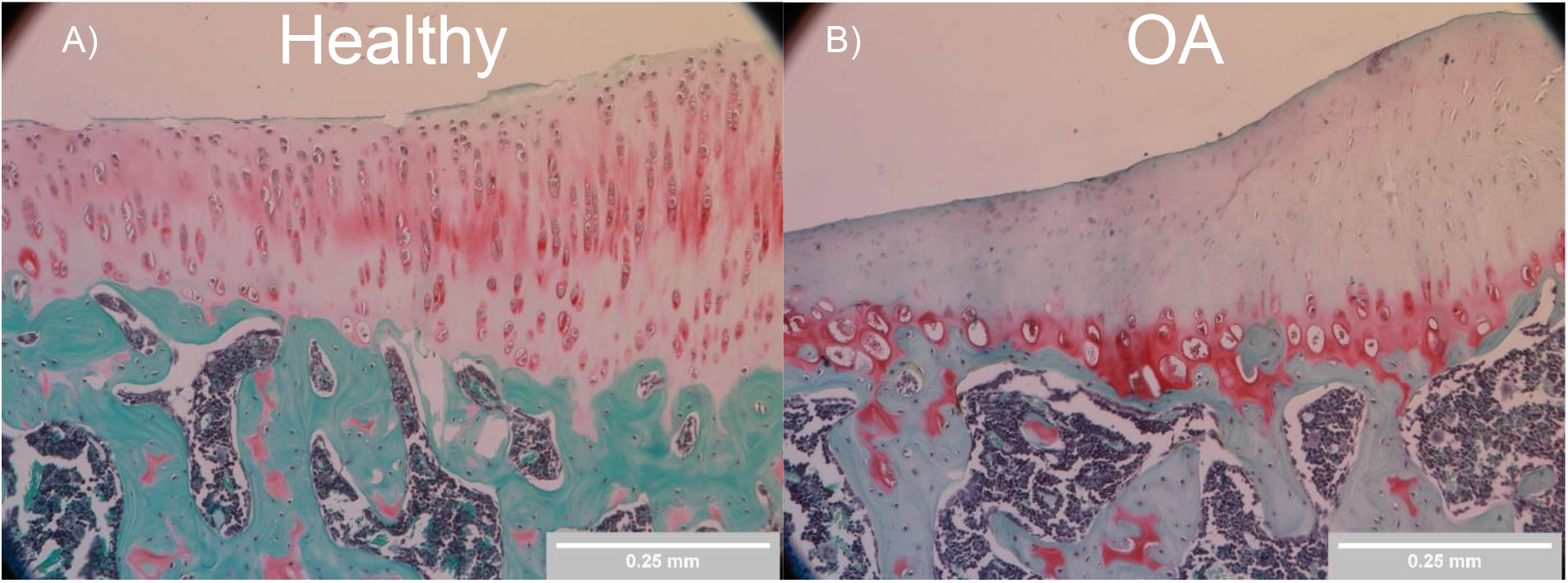
(column width). Histological staining of healthy and osteoarthritic tibiae. Coronal sections of the tibiae were stained with Safranin-O and Fast Green at 336 hr end point. A) CHP injected stifle joint demonstrates healthy cartilage. B) 21-days post-injection of 10 mg/mL MIA. There is an apparent loss of proteoglycans, loss of chondrocytes, and an increase in calcified cartilage at the osteochondral interface. Representative animal selected for each condition (n=4 per group).

## DISCUSSION

We examined the potential of CHPs to extend the retention time of imaging markers following IA injection, since CHP was previously demonstrated to bind to denatured collagens in joint tissues^[21,22]^. We and others demonstrated fluorescent CHP binding to joints of normal and diseased mice after tail vein injection. When CHP was delivered systemically via circulation, there was high CHP accumulation at the cartilage which was intensified by disease and injury.

Since local injection is a better drug delivery route for managing OA, and retention in the joint is a known issue for IA therapeutics used in OA treatment, we sought to determine if CHPs could bind to joint tissues for extended periods after IA injection. This was tested by in vivo imaging of rat knees after IA injection of fluorescently labeled CHPs (Fig. 1-3). Because of the high tissue penetration of near infrared light, NIRF was conjugated to active CHP and CHP with a scrambled sequence (sCHP), which has no affinity to collagen. We showed that CHPs substantially extend the retention time of NIRF imaging markers in both healthy and OA knee joints, with significant concentrations remaining at the final two-week imaging time point. By comparison to the scrambled sequence CHP, we showed that the enhanced retention of NIRF was due to CHP binding to denatured collagen, not the increase in molecular size. Healthy joints, in both humans and rats, experience a normal turnover of collagen during growth and remodeling that degrades collagen producing denatured collagen strands targeted by the CHPs.

The results for total radiant efficiency, AUC and AUMC indicate that CHPs bind to OA joints to a greater extent than healthy knee joints. Collagen turnover and the amount of denatured collagen is substantially increased in disease states such as OA, providing increased sites for CHP binding. As we expected, sCHPs did not bind to denatured collagen and were cleared from the joint rapidly as shown by the negligible NIRF intensities within the treated joints (Figures 1-3). However, the remaining PK parameters, *MRT, t*_1/2_, and *k*_e_, indicate that the NIRF-CHP localized in OA and healthy joints for a comparable period of time (Table 1). These results suggest that free or unbound CHPs are cleared from healthy and OA joints at similar rates, but bound CHPs are retained to a significantly greater extent in OA joints due to increased levels of denatured collagen. This implies that the bioavailability of the CHP is greater in the OA joints, which is promising for the future use with therapeutics.

After determining that CHPs were retained within the knee joint for the entire imaging period, we examined the specific CHP binding location within the joint tissue (Fig. 4). Using fluorescence imaging of the dissected joints, we learned that CHPs bind to multiple collagenous tissues within the knee joint, including cartilage, menisci, ligaments and synovial membrane. The highest fluorescence intensity was found on the central portion of the femoral and tibial cartilages, indicating CHP binding to the cruciate ligaments. This was to be expected given the high collagen content of ligaments, where CHPs can easily access and bind to the fibrous type I collagen in ligaments, as compared to accessing the dense matrix of type II collagen in cartilage. It is also possible that cutting ligaments during tissue isolation may have created a recoil in the ligament, which resulted in artificially bright fluorescence.

Finally, the development of OA was confirmed following injection of MIA into the rat knee joint using histological evaluation (Fig. 5). The changes observed during macroscopic evaluation of OA joints agree with what was previously reported in the literature^[24]^. With the development of OA following IA injection of MIA, the cartilage matrix showed characteristic changes similar to human OA, including loss of proteoglycans and hypocellularity through the cartilage thickness, as well as an increase in calcified cartilage at the osteochondral interface.

Other approaches have been used to extend the retention of therapeutics following IA injection. By simultaneously injecting hyaluronic acid (HA)-rhodamine and a thiolated HA-binding peptide, which targets type II collagen, Singh et al. demonstrated that the retention of hyaluronic acid (HA)-rhodamine could be extended^[10]^. However, this study only examined retention out to 72 hours and was only conducted in healthy animals. Another common method involves loading therapeutics into charged micro- or nano-particles that can interact with cartilage matrix. Although 10 um microparticles were retained in the joint^[9]^, there are potential complications of a foreign body response^[27,28]^ and a limited ability to penetrate into the cartilaginous matrix^[9,29]^. Rothenfluh and coworkers demonstrated that NPs functionalized with a ligand that binds to collagen II-alpha1 were retained in the joint 72-fold more at 48 hours post-injection compared to the NPs functionalized with scrambled sequence^[30]^. We believe that CHPs may address limitations of previous attempts to prolong therapeutic retention. First, our peptides directly target denatured collagen of many types (e.g. collagen type I in ligaments and collagen type II in cartilage), which are abundant throughout the joint and increased with the development of OA. Second, once bound, our peptides offer stable and long-lasting interactions with the collagen, providing the time necessary for therapeutic mechanisms of action to occur.

The MIA model used in this study is well suited for demonstrating CHP targeting of the OA condition, with the understanding that no OA animal model perfectly replicates all characteristics of human disease. Rats never experience closure of the epiphyseal plates, therefore cartilage and bone are constantly remodeling and forming, which is in contrast to humans where epiphyseal closure occurs following adolescence. Therefore, a higher rate of collagen turnover may be expected in rat models, as well as more accelerated changes to cartilage. Since our primary intent was to capture changes to the cartilage matrix, including loss of proteoglycans, loss of chondrocytes and cartilage calcification, as well as increased collagen turnover, the MIA model of OA was a reasonable choice. We also note that the MIA model is quite an aggressive model which develops rapidly. However, to demonstrate that CHPs have the ability to extend the retention time of IA injected compounds, the rapid pace only helps assert that CHPs can be retained even with the vicious nature of the disease progression. Further, by demonstrating that CHPs extend the retention time in healthy joints with lesser a degree of denatured collagen, there is still a possibility for extended retention in slower, spontaneous disease models such as that of primary OA in humans. In our study, we began the CHP injections at a time point that reflected changes in the joint that are representative of grade 2 OA, when there are subtle changes to cartilage and subchondral bone, but a distinct lack of cartilage delamination. This is a relevant stage since it reflects the point at which many patients become symptomatic and seek treatment. Finally, only healthy animals were used to examine CHP binding localization, leaving the CHP binding distribution in OA joints unknown. MIA leads to increased collagen damage in the joint, therefore increasing the extent to which CHPs bind. Consequently, we expect to see greater levels of CHP binding within each OA joint component compared to the healthy counterparts, while retaining similar binding distributions.

In summary, there is great potential to extend the retention time of therapeutics in the IA space of both healthy and OA joints using CHPs. Previously, we showed that CHPs can be conjugated to large biologic drugs such as the Fab region of Infliximab, and that the conjugates can be targeted to skeletal tissues via intravenous injection^[12]^. By conjugating CHPs to known disease-modifying or symptom-alleviating OA drugs, IA injected therapeutics could be retained in the joint space longer and at higher concentrations, allowing for improved efficacy, and longer intervals between treatments, ultimately slowing the rate of progression and delaying or eliminating the need for joint replacement. Further, CHP conjugation may allow use of low doses of drugs avoiding potential issues with off-target toxicity. Our study is the first step towards validating the capabilities of CHPs to extend the retention time of therapeutics for the treatment of OA.

## ACKNOWLEDGEMENTS

Financial support from NIH #R01AR071358 (JAW, SMY), NIH R21OD026618 (SMY), and NSF GRFP #1747505 (ENL) is gratefully acknowledged. We thank Mick Jurynec for helpful conversations, Perkin Elmer for assistance with interpreting well plate calibration, and Greg Stoddard for statistical guidance. IVIS imaging was performed at the University of Utah Preclinical Research Shared Resources Core. Mass spectrometry analyses of the synthesized NIRF-CHPs were performed at the Mass Spectrometry and Proteomics Core Facility at the University of Utah. Mass spectrometry equipment was obtained through a Shared Instrumentation Grant 1 S10 OD018210 01A1. SMY is a cofounder of 3Helix which commercializes CHP. All other authors have no competing interests to declare.

## REFERENCES

[1] Evans JT, Evans JP, Walker RW, et al. 2019. How long does a hip replacement last? A systematic review and meta-analysis of case series and national registry reports with more than 15 years of follow-up. Available from: http://www.thelancet.com

[2] Menkes CJ. 1994. Intraarticular treatment of osteoarthritis and guidelines to its assessment. J Rheumatol Suppl 41: 74–76.

[3] Rai MF and Pham CT. 2018. Intra-articular drug delivery systems for joint diseases. Current Opinion in Pharmacology 40: 67–73.

[4] Laurent TC and Fraser JRE. 1992. Hyaluronan. The FASEB Journal 6: 2397–2404.

[5] Simkin PA. 1995. Synovial perfusion and synovial fluid solutes. Annals of the Rheumatic Diseases 54: 424–428.

[6] Makarczyk MJ, Gao Q, He Y, et al. 2021. Current Models for Development of Disease-Modifying Osteoarthritis Drugs. Tissue Engineering - Part C: Methods 27: 124–138.

[7] Cai X, Yuan S, Zeng Y, et al. 2021. New Trends in Pharmacological Treatments for Osteoarthritis. Frontiers in Pharmacology 12.

[8] Ho MJ, Kim SR, Choi YW, and Kang MJ. 2019. Recent advances in intra-articular drug delivery systems to extend drug retention in joint. Journal of Pharmaceutical Investigation 49: 9–15.

[9] Pradal J, Maudens P, Gabay C, et al. 2016. Effect of particle size on the biodistribution of nano- and microparticles following intra-articular injection in mice. International Journal of Pharmaceutics 498: 119–129.

[10] Singh A, Corvelli M, Unterman SA, et al. 2014. Enhanced lubrication on tissue and biomaterial surfaces through peptide-mediated binding of hyaluronic acid. Nature Materials 13: 988–995.

[11] Hwang J, Huang Y, Burwell T, et al. 2017. In Situ Imaging of Tissue Remodeling with Collagen Hybridizing Peptides. ACS Nano 11: 9825–9835.

[12] Arlotta KJ, San BH, Mu HH, et al. 2020. Localization of Therapeutic Fab-CHP Conjugates to Sites of Denatured Collagen for the Treatment of Rheumatoid Arthritis. Bioconjugate Chemistry 31: 1960–1970.

[13] Li Y, Foss C, Summerfield D, et al. 2012. Targeting collagen strands by photo-triggered triple-helix hybridization. PNAS 109: 14767–14772.

[14] Li Y, Ho D, Meng H, et al. 2013. Direct detection of collagenous proteins by fluorescently labeled collagen mimetic peptides. Bioconjugate Chemistry 24: 9–16.

[15] San BH, Li Y, Tarbet EB, and Yu SM. 2016. Nanoparticle Assembly and Gelatin Binding Mediated by Triple Helical Collagen Mimetic Peptide. ACS Applied Materials and Interfaces 8: 19907–19915.

[16] Zitnay J, Li Y, Qin Z, et al. 2017. Molecular level detection and localization of mechanical damage in collagen enabled by collagen hybridizing peptides. Nature Communications 8.

[17] Zitnay J, Seob JG, Lin A, et al. 2020. Accumulation of collagen molecular unfolding is the mechanism of cyclic fatigue damage and failure in collagenous tissues. Science Advances 6.

[18] Lin A, Allan A, Zitnay J, et al. 2020. Collagen denaturation is initiated upon tissue yield in both positional and energy-storing tendons. Acta Biomaterialia 118: 153–160.

[19] Lin A, Zitnay J, Li Y, et al. 2019. Microplate assay for denatured collagen using collagen hybridizing peptides. Journal of Orthopaedic Research 37: 431–438.

[20] Zitnay J, Lin A, and Weiss JA. 2021. Tendons exhibit greater resistance to tissue and molecular-level damage with increasing strain rate during cyclic fatigue. Acta Biomaterialia 134: 435–442.

[21] Bennink L, Smith DJ, Foss CA, et al. 2017. High Serum Stability of Collagen Hybridizing Peptides and Their Fluorophore Conjugates. Molecular Pharmaceutics 14: 1906–1915.

[22] Bennink L, Li Y, Kim B, et al. 2018. Visualizing collagen proteolysis by peptide hybridization: From 3D cell culture to in vivo imaging. Biomaterials 183: 67–76.

[23] Dunshee LC, McDonough R, Price C, and Kiick KL. 2022. Retention of peptide-based vesicles in murine knee joints after intra-articular injection. Journal of Drug Delivery Science and Technology 74: 103532.

[24] Janusz MJ, Hookfin E, Heitmeyer S, et al. 2001. Moderation of iodoacetate-induced experimental osteoarthritis in rats by matrix metalloproteinase inhibitors. Osteoarthritis and Cartilage 9: 751– 760.

[25] Guzman RE, Evans MG, Bove S, et al. 2003. Mono-Iodoacetate-Induced Histologic Changes in Subchondral Bone and Articular Cartilage of Rat Femorotibial Joints: An Animal Model of Osteoarthritis. Toxicologic Pathology 31: 619–624.

[26] Bove SE, Calcaterra S, Brooker R, et al. 2003. Weight bearing as a measure of disease progression and efficacy of anti-inflammatory compounds in a model of monosodium iodoacetate-induced osteoarthritis. Osteoarthritis and Cartilage 11: 821–830.

[27] Pradal J, Zuluaga M, Maudens P, et al. 2015. Intra-articular bioactivity of a p38 MAPK inhibitor and development of an extended-release system. European Journal of Pharmaceutics and Biopharmaceutics 93: 110–117.

[28] Horisawa E, Kubota K, Tuboi I, et al. 2002. Size-Dependency of DL-Lactide/ Glycolide Copolymer Particulates for Intra-Articular Delivery System on Phagocytosis in Rat Synovium. Pharmaceutical research 19: 132–139.

[29] A. Maroudas. 1976. Transport of solutes through cartilage: permeability to large molecules. Journal of Anatomy 122: 335.

[30] Rothenfluh DA, Bermudez H, O’Neil CP, and Hubbell JA. 2008. Biofunctional polymer nanoparticles for intra-articular targeting and retention in cartilage. Nature Materials 7: 248–254.

